# AT vs. GC binding of protamine-template: A microscopic understanding through molecular dynamics and binding free energies

**DOI:** 10.1101/2025.04.17.649418

**Authors:** Sandip Mandal, Khadka B. Chhetri, Yun Hee Jang, Yves Lansac, Prabal K. Maiti

## Abstract

Protamine, an arginine-rich protein, compacts DNA more tightly than histones in somatic cells, yet its sequence-specific binding remains unclear. Using all-atom MD simulations with an arginine-rich short cationic peptide that mimics the protamine characteristics, we discovered distinct sequence preferences: the peptide binds preferentially to GC-rich sequences in the major groove and AT-rich sequences in the minor groove. Our structural analysis reveals that GC-rich binding induces significant DNA bending, narrowing the major groove and enhancing peptide interactions. In contrast, AT-rich minor grooves are more extended and electronegative, allowing better stereochemical fitting with planar and aromatic guanidinium side groups of arginine. However, thymine’s methyl group hinders major groove binding, favoring guanine. Thermodynamic free energy calculations, including MMGBSA, Jarzynski’s Equality, and Umbrella Sampling, confirm stronger peptide affinity for AT-rich motifs in the minor groove and GC-rich motifs in the major groove. Overall, this study will contribute to enhancing our understanding of DNA condensation and compaction in sperm cells based on protamine’s sequence-specific interactions.

## I. INTRODUCTION

Protein-DNA interactions are crucial for many biological functions, such as DNA replication, transcription, packing, and repair^1^. To understand these kinds of cellular processes, looking into the nature of protein-DNA complexes is crucial. The protein interacts with DNA in two different ways: first, recognizing the distinct chemical signatures of the DNA bases (base readouts); second, detecting DNA shape variations dependent on the sequence (known as shape readouts)^2^. The base readouts involve interaction between bases (A, T, G, and C) of the DNA and protein in the major and minor grooves, where proteins discriminate between the DNA bases through shape complementarity and electrostatic properties by forming several hydrogen bonds^2,3^. The DNA shape readout depends on the protein-induced DNA deformations^4,5^. Another important factor of specific protein–DNA recognition is protein flexibility^6–8^. The DNA-binding domains of the protein, which are highly specific, show more conformational changes when bound to DNA and greater flexibility when unbound^9^. Therefore, we must treat DNA and proteins as equal partners in order to fully understand the fundamental ideas governing protein-DNA recognition^10^. Hydrogen bonding (H-bonding) between proteins’ side chains and DNA bases determines the binding specificity of protein-DNA complexes^7^. The wider major groove is a suitable site for binding to relatively larger proteins, whereas the narrower minor groove is the binding site for smaller proteins like enzymes^11^. Most of the molecules that bind to the minor grooves are made of aromatic hydrocarbon rings^12^. DNA minor groove allows the molecule to fit into it by displacing water molecules out of the minor grooves^13–15^. The enzymes and transcription regulatory proteins mainly target the minor groove of DNA, which is a promising area for the development in therapeutics^16^.

Peptides interact with DNA through hydrophobic and hydrogen bond interactions in the major groove, while electrostatic interactions dominate at the phosphate backbone^17^. In the minor groove, the negatively charged sugar-phosphate backbones are closer together, making electrostatic interactions crucial for the protein binding^17^. Proteins fit tightly into both the major and minor grooves in DNA-protein complexes^18^, with sequence-specific changes in the DNA grooves are crucial for DNA-protein recognition^2,18,19^. While the major groove dimensions are consistent in both free and bound DNA, the minor groove dimension changes significantly upon binding^18,20^.

An X-ray diffraction analysis reveals more contacts in the minor groove of DNA than anticipated^21^. This strong interaction remains intriguing because the DNA minor groove is generally believed to be too narrow to allow protein components without causing significant energetic distortions^22^. According to Rohs et al.^18^, the minor groove’s electrostatic potential is affected by its shape, which proteins may be able to recognize. Poisson-Boltzmann calculations^18^ indicate that narrower minor grooves have a highly negative electrostatic potential, attracting basic residues like arginine. This electrostatic effect is especially pronounced in AT-rich regions, which often adopt very narrower minor grooves in free as well as bounded DNA structures^18^. This intriguing finding sheds light on how proteins might access DNA minor grooves, but it leaves the question of how the minor groove enlarges in many bound DNAs^18,20^. Crystallographic studies of DNA-protein complexes revealed that different structural configurations of the DNA backbone might be involved in minor groove binding processes that alter the groove dimension^21,23^. DNA-protein interactions are thus based on the intrinsic, sequence-dependent deformability of DNA. In protein-bound DNA, the stiffer AT-rich and flexible GC-rich segments are highly related with narrow and wider minor grooves, respectively^2,22^. DNA binding proteins identify such sequence-dependent DNA structures, where variation in DNA groove diameters is crucial^2^.

Protamines are basic proteins responsible for resulting in highly compacted and tightly coiled DNA in the sperm cells^24,25^. The genetic material is protected during spermatogenesis by this close DNA packing within the sperm head^26^. However, the protamine’s precise arrangement inside DNA-protamine complexes is still an unanswered question. According to the X-ray diffraction studies by Feughelman et al.,^27^, protamines are bound within the DNA minor groove. Raman spectroscopy on poly-arginine and salmon protamine^28^ and X-ray diffraction studies on arginine peptides^29^ have revealed that the protamines are located inside the major grooves. Recent molecular dynamics (MD) simulation studies show that protamine bound in the major groove utilizes the DNA backbones more effectively^30–32^. According to Mukherjee et al., the minor groove is too narrow for protamine binding, which causes severe DNA deformations, including unstacking and untwisting due to strong intercalative interaction between planar and aromatic guanidinium side group of arginine and AT base pairs^30^.

In order to figure out the unclear experimental/simulation questions from previous studies concerning protamine preference for DNA groove binding and its sequence dependence, we have conducted all-atom MD simulations using an arginine-rich short cationic peptide as a protamine-template. We have designed protamine-DNA complexes such that the short cationic peptide (mimic of protamine) gets bound to the major or minor groove of DNA during simulation. Our study focused on DNA-protamine complexes involving two 20 base paired oligonucleotides with sequences-poly(dAdT):poly(dAdT) and poly(dGdC):poly(dGdC). The peptide was designed to bind either to the minor or major groove. Comprehensive analyses, including hydrogen bonding, number of contacts, electrostatic interactions, and binding free energy calculations, reveal the sequence-dependent dynamics of protamine-DNA binding.

## II. COMPUTATIONAL DETAILS

### A. Model Building

The NAB^33^ code of AMBER20^34^ was used to prepare the twenty base-pair long double-stranded B-form DNA structures. Using the xleap module of the AMBER20, the short cationic peptide ACE-RRRSRRRS-NME was built, where R represents arginine (ARG), and S represents serine with a 3:1 arginine-to-serine ratio as protamine contains 60 to 80% arginine(R). This short cationic peptide (protamine-template) mimics the minimal properties of protamines effectively (see section S.1 in the supplementary information (SI) for a detailed description of protamine-template). The amide (NME) group (H_2_N−CH_3_) and acetyl (ACE) group (H_3_C=O) cap the N-and C-terminals, respectively, to prevent the terminal peptide chain interactions. We built DNA duplexes with two different DNA sequences: poly(dAdT):poly(dTdA), i.e., d(ATATATATATATATATATAT) and poly(dGdC):poly(dCdG), i.e., d(GCGCGCGCGCGCGCGCGCGC). To explore peptide-DNA binding preferences, we built four distinct peptide-DNA complexes: two for each DNA sequence (poly(dAdT):(dTdA) as d(AT) and poly(dGdC):(dCdG) as d(GC)), with the peptide positioned in either the major or minor groove (see Figure 1, upper panel). The interactions of DNA, as well as the native peptides, are represented by the Amber ff99SBildn force field^35^. Using the TIP3P water model^36,37^, each complexes were solvated, creating a 12 Å TIP3P water buffer around the structure. After adding the required quantity of Na^+^ (for DNA) and Cl^−^ (for peptide) ions to neutralize the system, additional amounts of Na^+^ and Cl^−^ ions were added to achieve a physiological salt concentration of 0.15 M. The ion interactions with water and solute atoms were studied using the Joung–Cheatham ion parameters set^38^. AMBER20^34^ simulation package was used for the unbiased MD simulation of the systems.

**FIG. 1.**
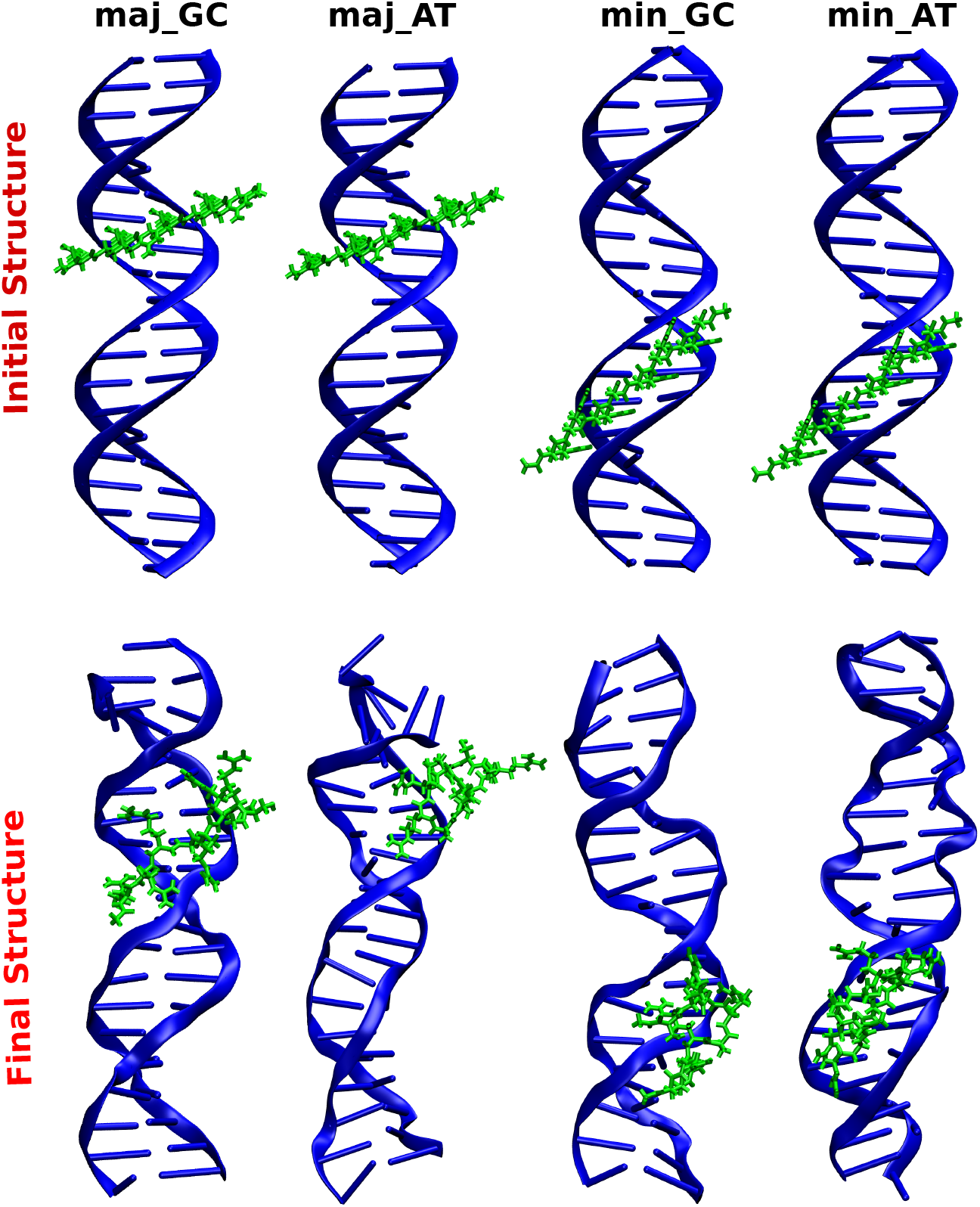
Instantaneous snapshots of the initial structures (upper panel) and the final structures(lower panel) from the 300 ns long MD simulation, i.e., after proper adsorption of the peptide in the DNA groove) of different complexes (DNA + peptide) in all-atom MD simulations. From left to right, the structures are: peptide bound to the major groove of d(GC) duplex, peptide bound to the major groove of d(AT) duplex, peptide bound to the minor groove of d(GC) duplex, and peptide bound to the minor groove of d(AT) duplex.

### B. Simualtion Details

We performed the energy minimization of systems to eliminate any unusual contacts that emerged during the system preparation, using 2,500 steps of steepest descent, followed by 2,500 steps using the conjugate gradient algorithm. A harmonic potential was applied with a spring constant of 500 kcal.mol^−1^Å^−2^ to all solute atoms to keep them fixed. The restraints were gradually reduced over five cycles, each involving 5000 steps of energy minimization, to reduce the constraint on the solute atoms to zero. This indicates that during the final 5000 steps of equilibration, the position restraint placed on the solute atoms was fully removed. The energy-minimized systems were then heated in four stages: with temperatures ranging from 10-50 K, from 50-100 K, from 100-200 K, and 200-300 K. Throughout the entire heating process, a harmonic restraint of 20 kcal.mol^−1^Å^−2^ was used to constrain the position of the peptide and DNA atoms.

To regulate the temperature, the Langevin thermostat^39,40^ was used with a coupling constant of 0.5 ps. Then, the systems were subjected to 5 ns NPT equilibration, where the pressure was kept at 1 atm using the Berendsen weak coupling method^41,42^ with a coupling constant of 0.5 ps. Subsequently, a 300 ns MD simulation with 2 fs integration time steps was performed in the NVT ensemble. The hydrogen bonds were constrained during simulation using the SHAKE algorithm^43^. We applied the Particle Mesh Ewald (PME) method^44^ to consider the electrostatic interactions. A cut-off distance of 10 Å was used for LJ interaction, and both the electrostatics and van der Waals (vdW) interactions terminated at that cut-off. Similar simulation strategies have been effectively used for several of our prior studies on DNA and DNA-based nanostructures^31,32,45–49^.

### C. Theory for PMF Calculations using Jarzynski’s Equality

Several advanced approaches can quantify the sequence-specific binding affinity of the arginine-rich cationic peptide to the DNA major and minor grooves, including Steered Molecular Dynamics (SMD), which uses non-equilibrium pulling work to measure binding affinity. Consider a thermally stable system in contact with a heat reservoir at a given temperature T. Let *λ* be a controllable macroscopic parameter (reaction coordinate) that alters the system’s state. If a system is driven isothermally from an initial equilibrium state A (consistent with *λ* = *λ*_*i*_) to B (consistent with *λ* = *λ*_*f*_) following a particular protocol *λ* (t): *λ*_*i*_ → *λ*_*f*_, then the work performed during this transition is given by-

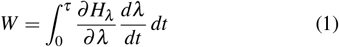

Here, *τ* denotes the transition time, indicating how quickly or slowly the transition process is completed, and *H*_*λ*_ represents how the system’s energy changes with the parameter *λ*. An ensemble of work values (W) can be obtained by repeating the above process multiple times.

Jarzynski first developed an equality connecting the non-equilibrium work performed using SMD and the equilibrium free energy difference (ΔF) between states A and B as given by:

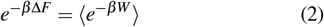

where ⟨·⟩ denotes the ensemble average over the work (*W*) values and 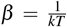. In this case, the ensemble average is calculated over multiple executions of the system’s non-equilibrium work performed during pulling simulations^50^. The work was computed for each pulling process as the integral of the force applied over the pulling distance to the target molecule’s center-of-mass (CoM) as –

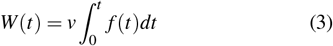

We employ a cumulant expansion based on a Gaussian distribution for approximation in order to address computational challenges arising from small work values in the exponential term. By using Jarzynski’s equality, we can calculate the profile of free energy along the reaction coordinate (*λ*). All other degrees of freedom are completely relaxed once the peptide reaches quasistatic equilibrium along this coordinate. The SMD simulations are utilized to sample the reaction pathway and calculate the Potential of Mean Force (PMF) using the stiff-spring approximation^51^-

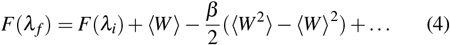

Through careful consideration of the computational complexity of non-equilibrium work measurements, we were able to capture the free energy landscape with this approach effectively.

### D. Theory for PMF calculation using Umbrella Sampling

In umbrella sampling (US), the binding free energy is calculated based on the population of the reaction coordinates. The final structures of the peptide-bound major and minor groove complexes generated by the all-atom MD simulations are stripped of the water molecules and the ions Na^+^ and Cl^−^, leaving only the peptide attached to the major and minor groove of the DNA. Each system was then solvated using a larger water box of dimension 100 × 60 × 100 Å^3^ along x,y, and z directions, respectively (the x-axis is the direction of pulling). It was then initially neutralized with counterions, and then more Na^+^ and Cl^−^ ions were added to get the desired physiological salt concentration of 0.15 M. A similar protocol described in the previous section was used for energy minimization, and then we used 10 ns NPT MD simulations to achieve equilibrium at 300 K.

Steered molecular dynamics (SMD) was performed to generate initial configurations for the umbrella sampling windows. The peptide was pulled away from the DNA binding site at a speed of 0.1 nm/ns in the x-direction using a spring force of 2000 kJ.mol^−1^.nm^−2^. During the pulling, the peptide was moved to 35 Å away from the DNA binding sites in steps of 1 Å. To prevent the DNA from being pulled along with the peptide during the pulling process, the heavier atoms of the DNA were restrained with a constant of 1000 kJ.mol.^−1^.nm^−2^. Constraints were imposed to ensure that the cationic peptide dissociates along a direction perpendicular to the DNA helix, avoiding rotation around the long axis of DNA. Umbrella sampling was employed to improve the sampling of the energetically unfavorable regions, allowing to compute the potential of mean force (PMF) by applying a harmonic bias potential of force constant of 2000 kJ.mol^−1^.nm^−2^ to the peptide’s center of mass during 30 ns equilibrium MD simulation in each window. As a result, the overall simulation duration for the US run was 1.05 *µ*s for each system. This sample size was sufficient to ensure that the PMF profile has converged. The PMF profiles for peptide-DNA dissociation were generated using the WHAM code implemented in GROMACS^52^, capturing the free energy landscape along the dissociation pathway.

## III. RESULTS AND DISCUSSION

### A. Structural Stability and Energetics

We begin by analyzing the effects of conformational stability and energetic contributions to sequence-dependent binding affinity from the unbiased simulation trajectory. Figure 1 (lower panel) shows the final conformations of peptide-adsorbed DNA complexes after 300 ns long unbiased MD simulations. The final DNA conformations retain their B-form, as confirmed by helical parameters calculated using 3DNA codes(see Table 2, in SI). The structural fluctuations of the peptide during 300 ns simulation are assessed by its RMSD values. The structure’s RMSD with respect to its reference structure at time *t* is provided by: 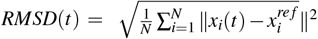, where {*x*_*i*_(*t*)} are the N atoms’ co-ordinates in the structure, while the coordinates of the initially energy-minimized structure are 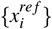. Among the different binding geometries, the configuration with the smallest RMSD is considered to be the most stable and potent binding configuration. It should be mentioned that the measures of different physical quantities presented throughout this manuscript are the averages taken over the last 50 ns out of 300 ns long MD trajectories. To quantify structural fluctuations at longer timescales, we have extended the simulations for minor groove binding of poly(AT) and major groove binding for poly(GC) up to 1 *µ*s and observed structural stability and similar binding affinity as 300 ns (see Section S.2 in SI for detailed analysis of structural properties and energetics from the 1 *µ*s extended simulations.

Figure 2A shows the RMSD of all the non-hydrogen atoms of peptide (protamine-template) relative to their initially minimized structures, which bind to the major and minor grooves. In the case of major groove binding, the peptide has a lower RMSD of 5.02 ± 0.29 Å for d(GC) and a higher RMSD of 8.09 ± 0.23 Å for d(AT). In minor groove binding, the peptide has a lower RMSD value of 6.43 ± 0.24 Å for d(GC) and a higher RMSD value of 7.99 ± 0.20 Å for d(AT). The lower stable RMSD value for d(GC) in the major groove is due to stronger hydrogen bonding and electrostatics. In contrast, for d(AT), the peptide has higher fluctuations with respect to simulation times, due to the steric hindrance by thymine(T) methyl groups (see Figure S3.A, in SI). In minor grooves, DNA-peptide binding is facilitated by stacking interaction between the guanidinium side groups of the ARG and d(AT) bases, along with the electrostatic interaction between the protamine ARGs and DNA phosphates. As a result, the peptide has a higher and stable RMSD value in the d(AT) minor groove over the last 200 ns of the production run as shown in Figure S3.B, SI. In contrast, for minor groove binding of d(GC), the peptide is found to adhere to the backbone of DNA rather than being fitted inside the groove, which might explain the lower but fluctuating RMSD. Although smaller RMSD value peaks indicate a stable configuration, due to stacking-induced deformation, the RMSD value for the peptide binding at the d(AT) minor groove initially increases and then stabilizes for the last 200 ns (Figure S3.B). The RMSD analysis of the complex (peptide + DNA) also reveals similar trends (see Figure S4, SI): for major groove binding, the complex is more stable with d(GC) and less with d(AT); for minor groove binding, the stability is reversed, with d(AT) being more stable than d(GC).

**FIG. 2.**
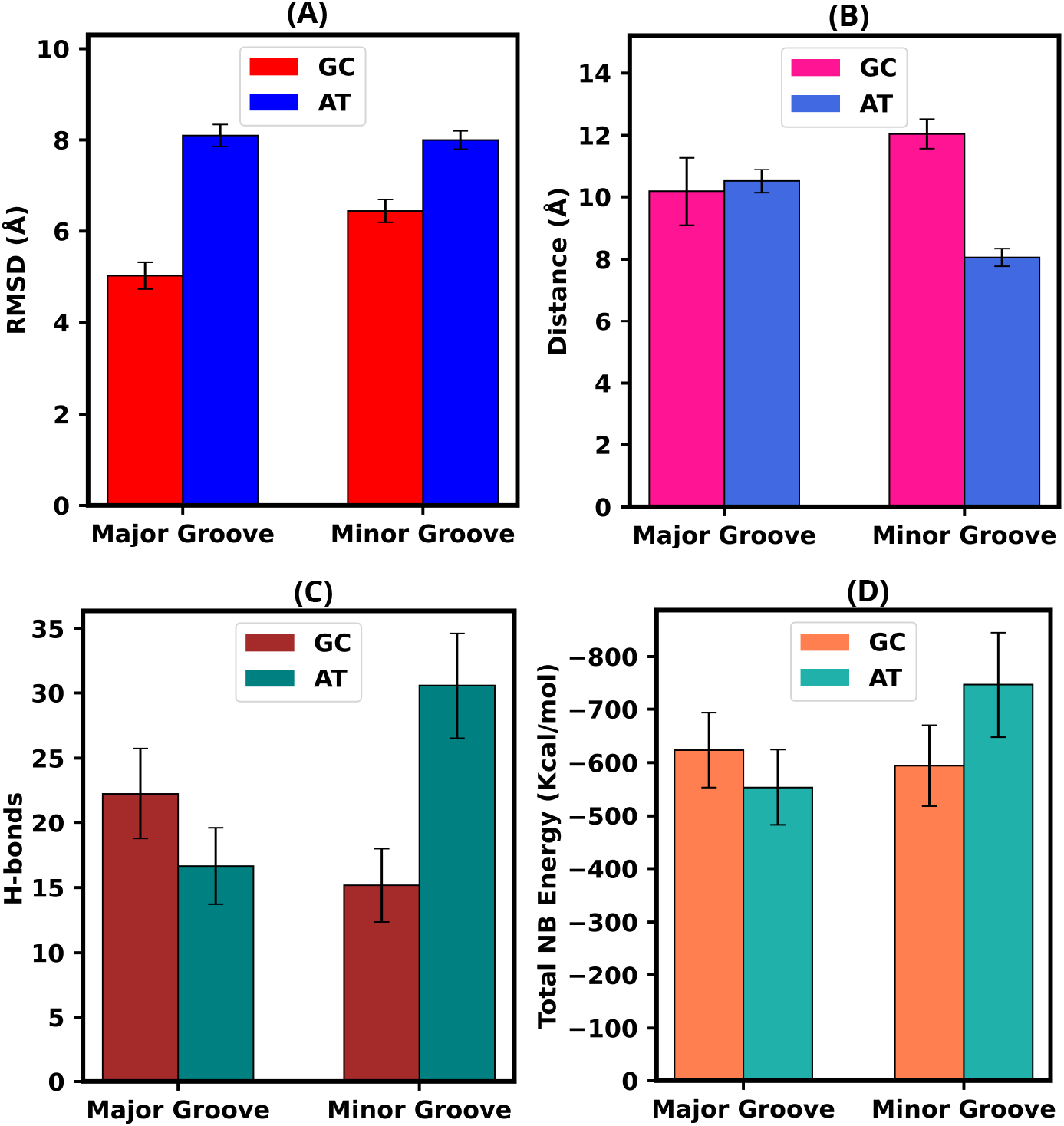
Histograms plots for the **[A]** Peptide’s RMSD of non-hydrogen atoms, **[B]** COM Distance between Peptide and DNA binding regions/residues as determined from distance-based contact maps, **[C]** Total non-bonded interaction energy (vdW+elec), and **[D]** Number of hydrogen bonds (H-bonds), for the major and minor grooves binding of the peptide to the poly(GC) and poly(AT) using the last 50 ns of the 300 ns long unbiased MD trajectory.

The peptide that fits well into the DNA groove remains closely embedded, resulting in a shorter distance between the peptide’s center of mass (COM) and the DNA binding site’s COM COM of DNA residues selected based on a distance-based contact map). For major groove binding, the COM distance is larger for the d(AT) duplex than for the d(GC) duplex (see Figure 2B). The peptide’s COM distance differs by DNA duplex sequences in the major and minor grooves: the average COM distance in the major groove is 10.19 ± 1.09 Å for d(GC) and 10.52 ± 0.37 Å for d(AT). The peptide is more deeply embedded in the major groove of d(GC) than in d(AT). However, for minor groove binding, the trend reverses as the peptide stays closer to d(AT), with a COM distance of 8.05 ± 0.28 Å, as compared to 12.05 ± 0.48 Å for d(GC) (Figure 2B). These findings suggest that binding is stronger in the major groove of d(GC) and in the minor groove of d(AT) due to closer proximity of the peptide.

The radius of gyration (*R*_*g*_) represents the root-mean-square distance between a polymer’s segments and its center of mass (COM). For proteins, *R*_*g*_ represents the distribution of atoms around their axis, with higher values for linear shapes and lower values for tightly packed, globular *α*/*β*-proteins^53^. The time evolution of *R*_*g*_ for peptide binding to the major and minor grooves of the DNA duplexes is shown in Figure S5.A and B in SI. The peptide’s *R*_*g*_ for major groove binding is greater for the d(GC) duplex (8.51 ± 0.32 Å) than for the d(AT) duplex (6.58 ± 0.11 Å), suggesting that in the d(AT) duplex, peptide has tighter packing and the d(GC) duplex has more extended conformation. In minor groove binding, the peptide has a slightly smaller *R*_*g*_ with the d(GC) duplex (6.77 ± 0.17 Å) than the d(AT) duplex (6.88 ± 0.24 Å) for the last 50 ns of the production run, indicating a more extended conformation in d(AT). This extended geometry improves peptide-DNA interactions by increasing the number of contacts with d(AT) bases, improving van der Waals and hydrophobic interactions, and providing a better stereochemical fitting in the narrow minor groove due to effective stacking interactions^30^. Thus, the *R*_*g*_ analysis indicates higher peptide binding affinity in the major groove of d(GC) and in the minor groove of d(AT). The DNA+peptide complex also follows the same trend (Figure S6.A and B, SI)

Numerous interactions, such as hydrogen bonding, cation*‐π*, stacking interactions, and water-mediated effects, have been investigated in relation to the contacts between DNA and proteins^54^. The primary mechanism of DNA-protein interactions is based on hydrogen bonds (H-bonds), which are essential for the protein’s ability to read the DNA target directly. The recognition of specific DNA sequences by proteins depends on the interaction between the hydrogen-bond donors and the acceptors of the DNA base pairs, as well as the protein’s amino acids that fit into the grooves^55^. We evaluated the H-bonds between the peptide and DNA using the Hydrogen Bond plugin from the VMD^56^ software. To ensure accurate measurements, the distance cut-off was set at 3.5 Å and the angle cut-off at 120^*o*^, following IUPAC recommendations^57^. For the major groove binding, the peptide forms more H-bonds with the d(GC) duplex than with the d(AT) duplex (Figure 2C). The average number of H-bonds made by the peptide with major groove d(GC) is 22.25 ± 3.48, and this number is 16.25 ± 2.93 when bound to d(AT) duplex. Whereas, for the minor groove binding, the peptide forms more H-bonds with the d(AT) duplex than with the d(GC) duplex (Figure 2C). The average number of H-bonds made by the peptide with the minor groove of d(GC) is 15.17 ± 2.80, and the number is 30.56 ± 4.05 when bound to the minor groove of d(AT) duplex. Consequently, it is evident from the number of H-bond counts that the cationic peptide binding favors the AT-rich motif in minor grooves and the GC-rich motif in major grooves.

Also, we have computed the number of contacts made by peptide with DNA using the CPPTRAJ^58^ tool provided with AMBER20 software^34^, taking a distance cut-off of 3.5 Å. The number of contacts made by the peptide with DNA duplexes also shows a similar trend as that of the number of H-bonds made by the peptide with the DNA duplexes for the major (Figure S7.A,SI) and minor groove (Figure S7.B, SI) bindings, respectively. In major groove binding, the peptide makes more contact with the d(GC) than with d(AT). The average number of contacts made by the peptide with d(GC) is 96.09 ± 12.08 and the number is 83.21 ± 11.0, when bound to the major grooves of d(AT). For the case of minor groove binding, the number of contacts formed by the peptide is relatively higher with d(AT) than with d(GC). The average number of contacts made by the peptide with the minor grooves of d(GC) is 78.29 ± 10.11, and the value is 139.47 ± 13.14 when bound to the minor grooves of d(AT). These results imply that the peptide prefers GC-rich motifs for major groove binding, and AT-rich motifs for minor groove binding. The relatively more significant number of H-bonds and other contacts made by the peptide in the minor groove than in the major groove should be associated with the narrow dimension and stacking interaction between the planar guanidinium side group and DNA bases at the minor groove^30^. In DNA, the minor groove containing AT-rich sequences, particularly A-tracts, is much narrower and more electronegative than the others^18,59^. Figure 3A-B clearly shows the higher binding preference for major grooves of d(GC) and minor grooves of d(AT), which are dominant among the four complexes and also higher contact surface area (CSA) for minor groove of d(AT) as compared to major groove of d(GC) due to intercalation (see Figure S.8, SI).

**FIG. 3.**
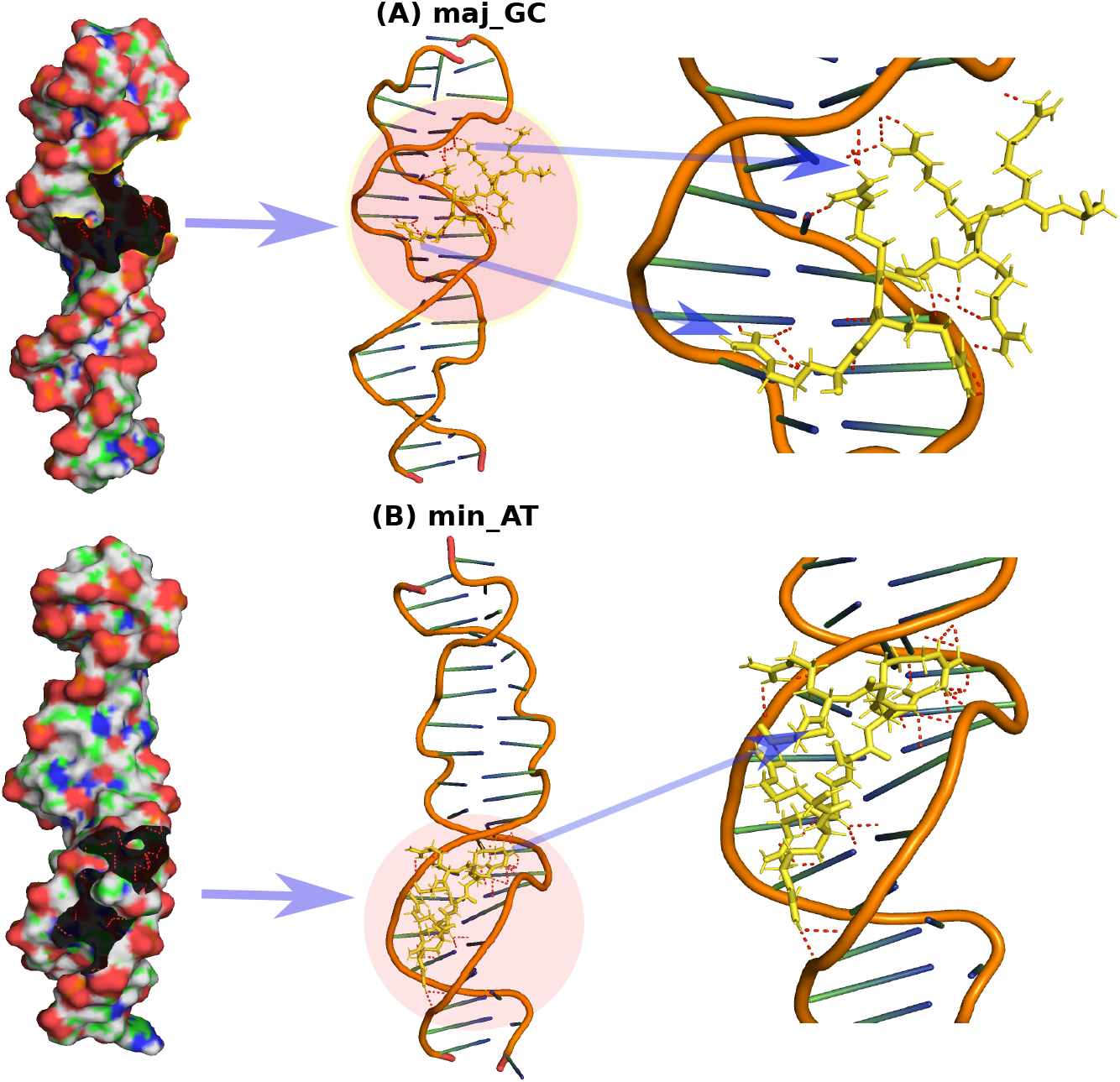
Instantaneous snapshots from the last frame of 300 ns MD simulation for major and minor groove binding of peptide with polar contacts. Since the major groove of d(GC) and minor groove of d(AT) show the most preferred binding affinity, we present only the two important DNA-peptide complexes out of the four. **[A]** The left figure is just a surface representation to show the peptide binding location in the major groove of d(GC), the middle and right figures indicate the binding of peptide with the formation of polar contacts, mostly with the backbones **[B]** The left, middle and right figures all together represent tight binding of the peptide in the minor groove of d(AT) with the formation of more polar contacts mostly with the DNA bases, which confirms stacking interaction of planar guanidinium side groups of arginine in d(AT) minor groove.

To quantify how the short cationic peptide interacts with DNA through various mechanisms, including electrostatic, van der Waals (vdW), and solvation forces. The vdW and electrostatic interactions between functional groups and mobile ions and physiological environmental effects are all energetic contributors in protein–DNA complexes^60,61^. The vdW attractions from the contact surfaces of the components are used to assess the role of shape complementarity in binding in macromolecular complexes^61^. Although several other interactions are involved in DNA-protein interactions, electrostatic interactions play a crucial role in protein-DNA binding^62^. We computed the NB interaction energies (vdW and electrostatic) between the peptide and the DNA using the NAMD Energy (MDEnergy) module^63^ interfaced with VMD software. The total Non-bonded interaction energies between the peptide and DNA are shown in Figure 2D. Figure S9.A-B and S.10.A-B in SI show energies to elucidate the vdW and electrostatic contributions to the total NB energy with respect to their variations with simulation time. The interaction is more favorable when the peptide binds to the major groove of the d(GC) and the minor groove of d(AT). Total NB energy is more dominant for the minor groove of the d(AT) than d(GC). For the major groove binding, the total NB energy is-623.39 ± 70.17 kcal.mol^-1^ for d(GC), and the value is-553.16 ± 71.17 kcal.mol^-1^ for d(AT). while in the case of minor groove binding, the total NB energy is-594.14 ± 75.82 kcal.mol^-1^ corresponding to d(GC) and the value is-746.12 ± 98.48 kcal.mol^-1^ for the d(AT). The AT-rich motif is preferred in the minor groove, whereas the GC-rich motif is favored for major groove binding.

The van der Waals (vdW) energy is more favorable to the major groove of the d(GC) than d(AT). In minor groove binding, vdW energy is more favorable for the d(AT) than d(GC). During major groove binding, the vdW energy is-46.25± 8.65 kcal.mol^-1^ when the peptide is bound to the major groove of the d(GC) duplex, and the value is-41.01 ± 7.46 kcal.mol^-1^ for d(AT). During minor groove binding, the vdW energy is - 36.58 ± 11.49 kcal.mol^-1^ when the peptide is bound to the minor groove of the d(GC), and the value is-66.32 ± 8.04 kcal.mol^-1^ when the peptide is bound to d(AT) duplex’s minor groove. Considering vdW energies only, these results suggest that the arginine-rich cationic peptide prefers GC-rich motifs for major grooves and AT-rich motifs for minor groove binding.

Electrostatic energy is more dominant when the peptide binds to the d(GC) major groove than d(AT). In contrast, electrostatic energy is more favorable in the d(AT) minor groove than d(GC). During major groove binding, the electrostatic energy is-577.14 ± 66.75 kcal.mol^-1^ when the peptide is bound to the major groove of the d(GC), and the value is - 512.15 ± 69.69 kcal.mol^-1^ for d(AT). For minor groove binding, the electrostatic energy is-557.54 ± 69.57 kcal.mol^-1^ for d(GC), and the value is-679.79 ± 93.84 kcal.mol^-1^ for the d(AT) duplex. Therefore, the NB electrostatic energy values strongly support our hypothesis that the arginine-rich cationic peptide favors GC-rich motifs for major groove binding due to the more bending angle of the DNA and extended conformations of peptide, helping to form more H-bonds and contacts. The peptide prefers AT-rich motifs for minor groove binding due to intercalative/stacking interactions (as shown in Figure 4.A-D through distance-based contact map and zoomed view of intercalation by identifying the residues involved) resulting in the highest electrostatic energy among all four complexes. These are the scenarios seen in other analyzed properties in the earlier sections. However, in the energetics, other polar and non-polar interactions are also related to the solvation-free energy. These results add up to decide the binding preference, which is discussed in the MMGBSA section next to it.

**FIG. 4.**
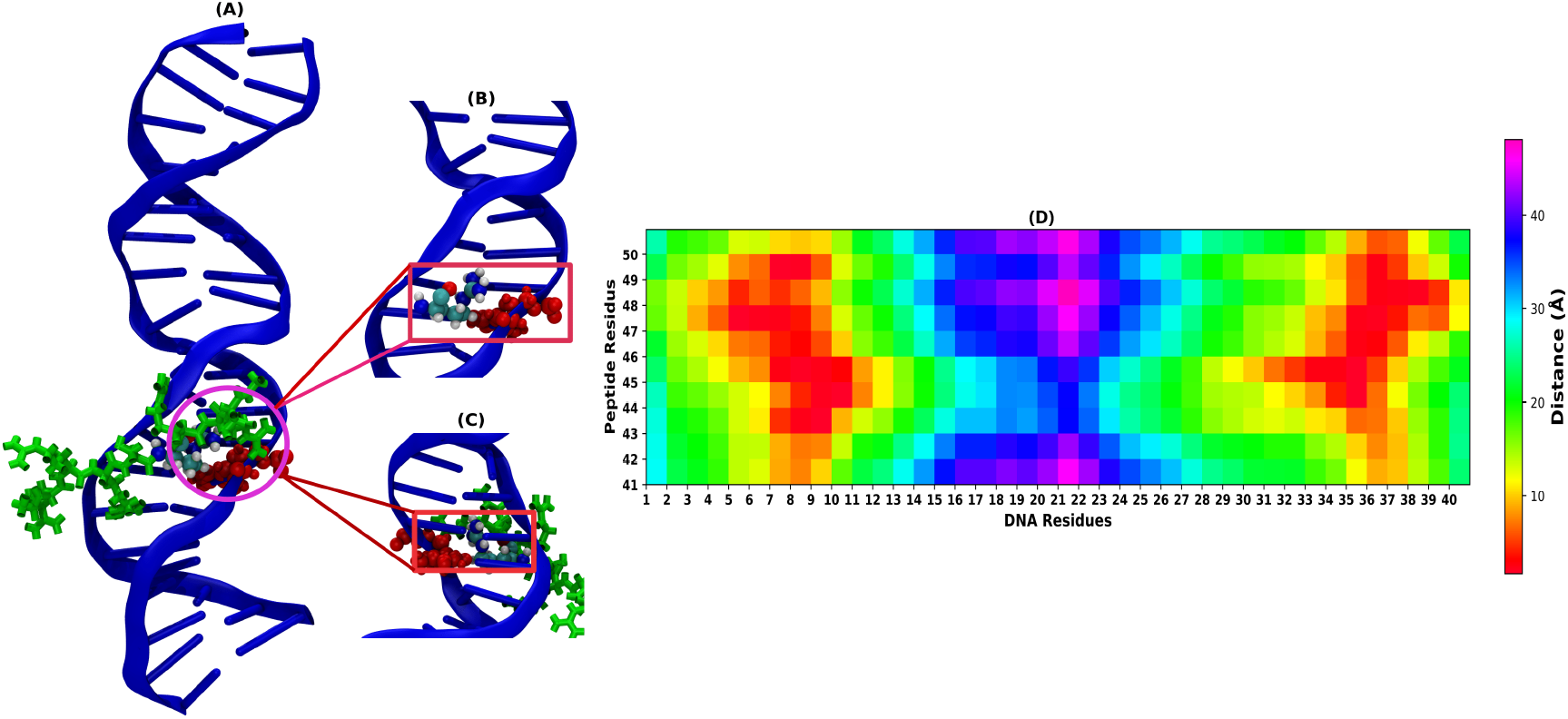
Representation of intercalation for ARG’s planner guanidinium groups to the minor grooves of AT-rich base pairs from the 1 *µ*s MD trajectory. (B) Zoomed front view of the guanidinium and minor groove’s AT base pair interaction, (C) Zoomed backside view of the stacking interaction. (D) Distance-based contact map between peptide residues (y-axis) and AT-rich DNA residues (x-axis), where the peptide is placed near the minor grooves of DNA.

### B. Binding Free Energy calculations from MMGBSA Approch

Our binding free energy calculation from the unbiased trajectory using molecular mechanics based generalized Born surface area (MMGBSA) approach, which is implemented in the MMPBSA.py^64^ module of AMBER20, shows that the enthalpic (ΔH) contribution for protamine binding to DNA major and minor groves decreases in the following order: min_AT *>>* maj_GC. It suggests that the arginine-rich cationic peptide tightly binds in the major groove of d(GC) and in the minor grooves d(AT) enthalpically (ΔH), with the electrostatic contact between the peptide’s positive arginine residues and the nucleotides’ negative phosphates controlling the associations. Due to the tight binding, peptides become less mobile, resulting in a more negative entropic change TΔS, as listed in Table I. The major groove of d(GC) has a more negative entropy value than the minor groove of d(AT), which could be related to the minor groove’s structural distortion due to the stacking interaction with the planar guanidinium side group of the peptide. In our work also, we observed solvation free energy difference (300 kcal/mol) between the major and minor grooves of AT- and GC-rich DNA arises due to distinct interaction strengths, similar effect was previously observed by Ghosh et al.^65^. The solvation-free energy (Δ*G*_*sol*_) contributes negatively to the peptide binding in both major and minor grooves, but it is more prominent in the minor grooves. This is because entering the DNA grooves requires the peptide to replace water molecules from the grooves, which is more difficult in the narrower minor groove. Therefore, due to the presence of water hydration, the peptide will lose a significant amount of H-bonds, and electrostatic interaction will be less favorable; therefore, the more positive solvation energy indicates that the peptide is not favorable for binding. Consequently, the solvation-free energy hindrance has been compensated through the stronger electrostatic interaction. Overall, the free energy upon peptide binding (ΔG = ΔH - TΔS) is most favorable for the peptide bound to the minor groove of d(AT), followed by the major groove of d(GC) as shown in Figure 5. All the energy component values from MMGBSA calculations are listed in Table I.

**TABLE I.**
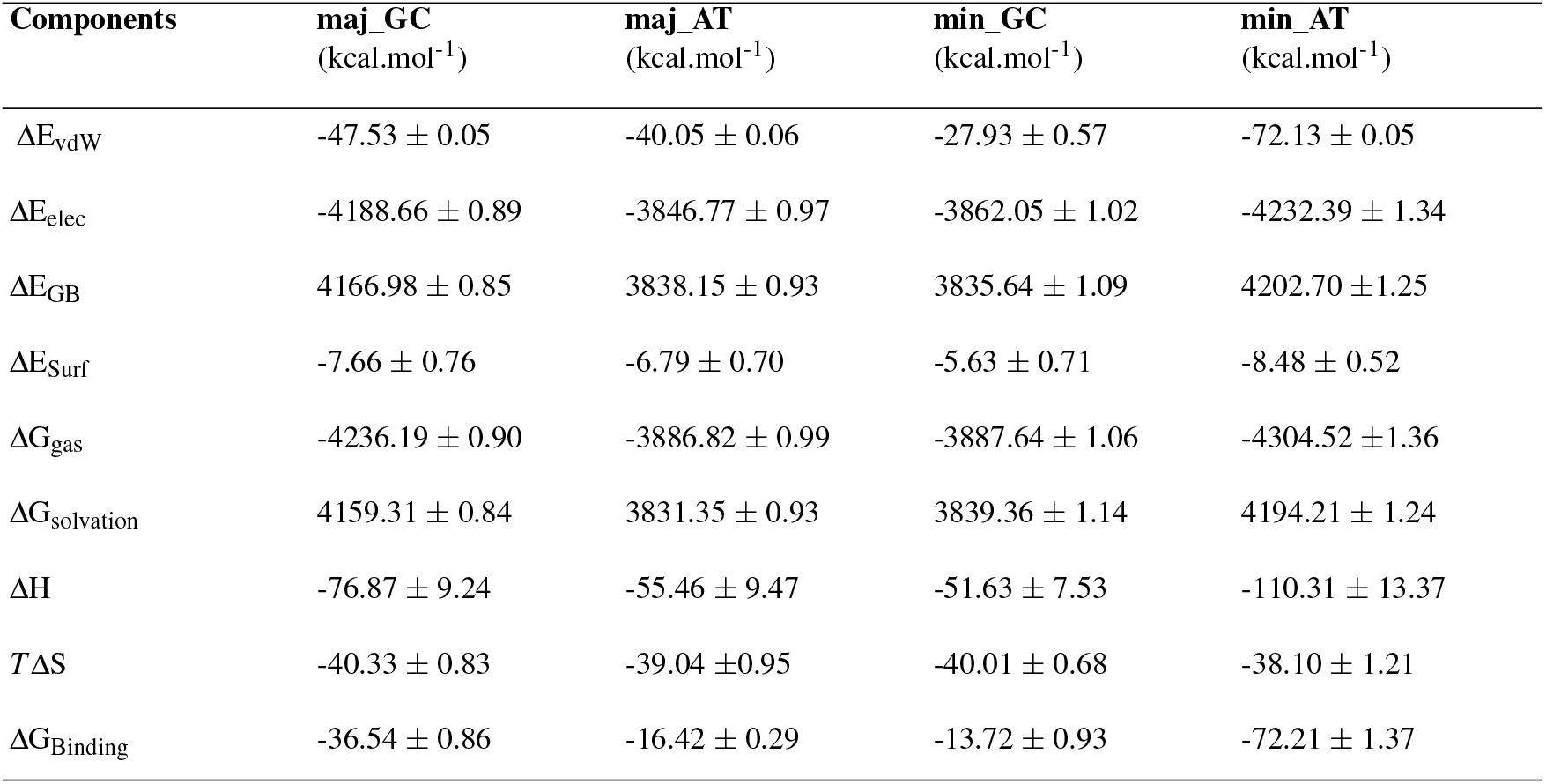
Binding free energy values from different energy components of the peptide’s binding to the major and minor grooves of d(GC) and d(AT), using the last 100 ns of the 300 ns long unbiased MD trajectory utilizing the MMGBSA methodology.

**FIG. 5.**
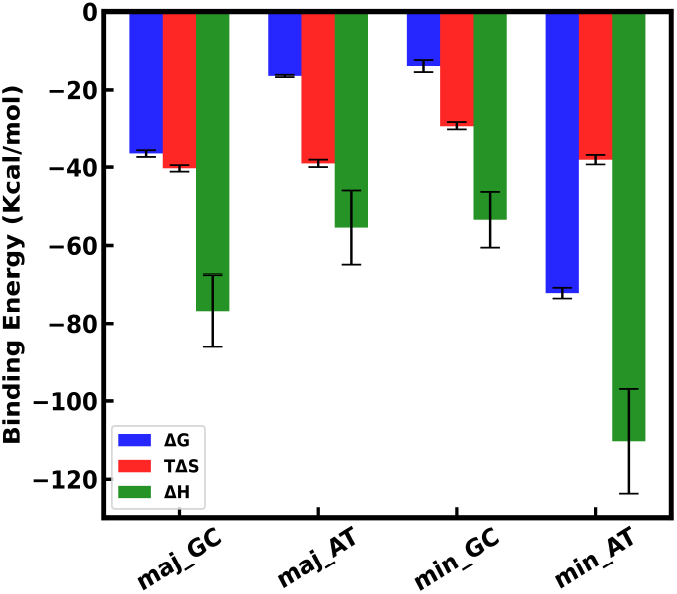
The binding free energies (ΔG = ΔH - TΔS) between the peptide and the DNA grooves using MMGBSA method from the last 100 ns of the 300 ns long MD simulation trajectory. The details of the different energy component values are listed in the Table I.

### C. Binding free energy from Jarzynski Equality

In this section, we have discussed results from the non-equilibrium SMD simulation trajectories. Jarzynski Equality (JE) provides a theoretical method for calculating the free energy difference by exponentially averaging the non-equilibrium works performed between bound and free states. Therefore, we employed JE to compute the potential of mean force (PMF) by pulling the cationic peptide away from the binding regions of the major and minor groves of DNA. The non-equilibrium work (equation 1,2,3) performed for moving the peptide in a specific direction (x-axis) over 10 trajectories at a very slow speed of 0.1 nm/ns are shown in Figure 6(A, B, D, and E) with a spring force constant of 2000 kJ mol^−1^ nm^−2^. However, significant variations in the work that deviates from the Gaussian distribution lead to poorly converged PMFs using only the first two cumulants of the work distributions (equation 4). Hence, we used the exponential averaging form of Jarzynski’s equality as provided by equation 2 to improve the outcomes. The unbound(free) state corresponds to the flattening of the PMF beyond a distance of 30 Å (3 nm). Using JE, calculated binding free energy values are Δ*G*_*GC*_ = 50.30 ± 0.94 kcal.mol^-1^ and Δ*G*_*AT*_ = 40.80 ± 3.56 kcal.mol^-1^, at the major groove, with the energy minima’s (*λ* _min) at 7.20 Å and 6.30 Å respectively. Similarly, the binding free energy values are Δ*G*_*GC*_ = 2.58 ± 0.01 kcal.mol^-1^ and Δ*G*_*AT*_ = 8.70 ± 0.09 kcal.mol^-1^ for the minor groove binding with the energy minima’s (*λ* _min) at 13.11 Å and 8.89 Å respectively. This indicates that d(GC) is preferred for major groove and d(AT) is more favored for minor groove binding. Interestingly, Figure 6(F) reveals proof of a strong stacking interaction between d(AT) and planar guanidinium side groups, where the peptide is embedded deep inside the minor groove of d(AT) (*<*8 Å). A second interaction is observed between the DNA backbone of d(AT) and the peptide near 13 Å, which is also the interaction region between the peptide and the minor groove of d(GC), as shown by the grey-shaded region.

**FIG. 6.**
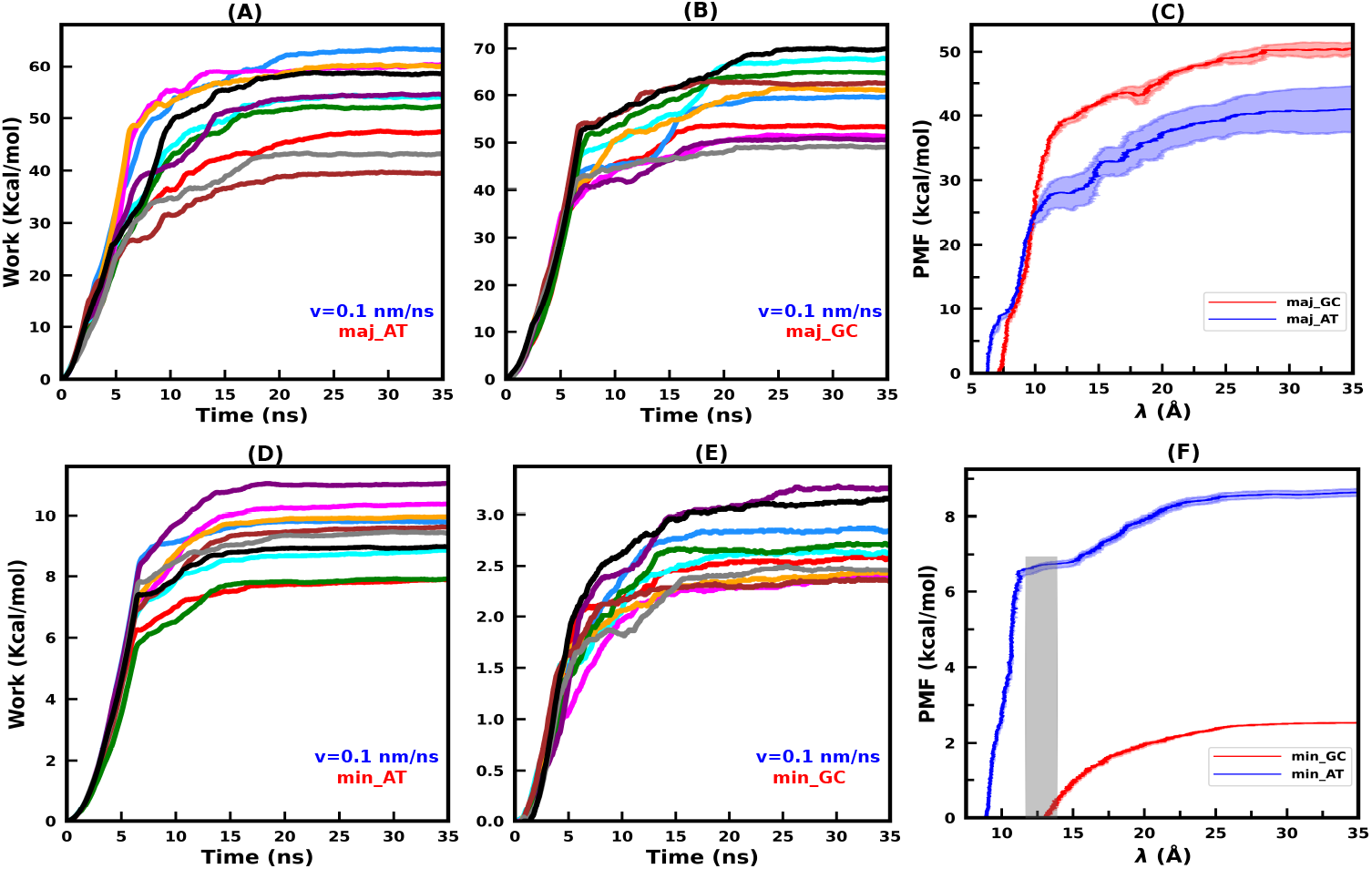
PMF and work profiles for slowly pulling the cationic peptides from major and minor grooves of DNA utilizing the SMD and Jarzynski’s equality for all 10 SMD trajectories. **[A, B]** shows the non-equilibrium work profiles of the peptide dissociation from the major grooves of d(AT) and d(GC). **[C]** represents the PMF profile for major grooves of d(GC) and d(AT) using JE with red and blue colors. **[D, E]** display work profiles of peptide dissociation from the minor grooves of d(AT) and d(GC) for 10 SMD trajectories with a speed of 0.1 nm/ns. **[F]** Shows PMF profiles in red and blue colors for d(GC) and d(AT) using JE for 10 SMD trajectories. The gray shaded bar represents the region of the second interaction that the peptide experiences with the DNA backbone (in the absence of stacking, *>*10 Å) in the d(AT) minor groove. The red and blue shaded regions represent the error bars, calculated based on the ensemble average of the last 8,9 and 10 trajectories.

We hypothesize that when we use JE to analyze the pulling trajectories and generate the PMF profiles, there is a significant restriction on the peptide’s rotation, indicating that SMD did not provide enough sampling to capture the peptide’s rotational degrees of freedom, as reported in the literature also for DNA-protein interaction^51^. This discrepancy implies that the peptide’s free energetics were affected, resulting in an overestimation of the absolute binding free energy Δ*G*_*b*_ for the major groove of d(GC) and d(AT). On the other hand, in the minor groove, the peptide’s structure becomes more compact due to the fitting in the narrower minor groove, which results in an underestimation of binding free energies. To overcome these limitations, we use umbrella sampling to improve sampling on the peptide’s rotational degrees of freedom, as discussed in detail in section S.9 in SI about the peptide’s rotational freedom in both cases. Overall, while JE over/underestimates the binding free energy values for major/minor grooves, it does provide the correct sequence-dependent trends for binding of the peptide to the major and minor groove of the DNA, as demonstrated in previous parts.

### D. Potential of mean force (PMF) from US

Figure 7 depicts the dissociation free energy profiles (PMF, *F*(*ζ*)) obtained from umbrella sampling simulations for peptide-bound DNA complexes in major and minor grooves. Each PMF *F*(*ζ*) profile features a global energy minimum at the location (*ζ*_*min*_), indicating the preferential binding distance (bound state). Beyond this global energy minimum, the PMF increases until it reaches a plateau where the DNA-peptide interaction is practically eliminated (unbound/free state). The free energy barriers between the bound and unbound states are higher for d(GC) compared to d(AT) in the major groove, while exactly the opposite trend is observed in the minor groove, consistent with all of our prior observations. For the major groove binding, the PMF profiles show well depths of-21.99 kcal.mol^-1^ for d(GC) and-17.83 kcal.mol^-1^ for d(AT), respectively, for the major groove binding. while for the minor groove binding, the well depths are-19.02 kcal.mol^-1^ for d(AT) and-13.23 kcal.mol^-1^ for d(GC). When the peptide does not experience the stacking interaction anymore (*<* 10Å) between the minor groove d(AT) base pairs and the planar guanidinium side group of ARG, it interacts again with the DNA backbone (*>* 10Å), causing a small local minima in the minor groove PMF profile of d(AT) (Figure 7B, red curve). We have also calculated changes in helical parameters (groove widths, bending angle, and conformational changes) at the bound/unbound states of the peptide and briefly discussed in Section S.8 in SI.

**FIG. 7.**
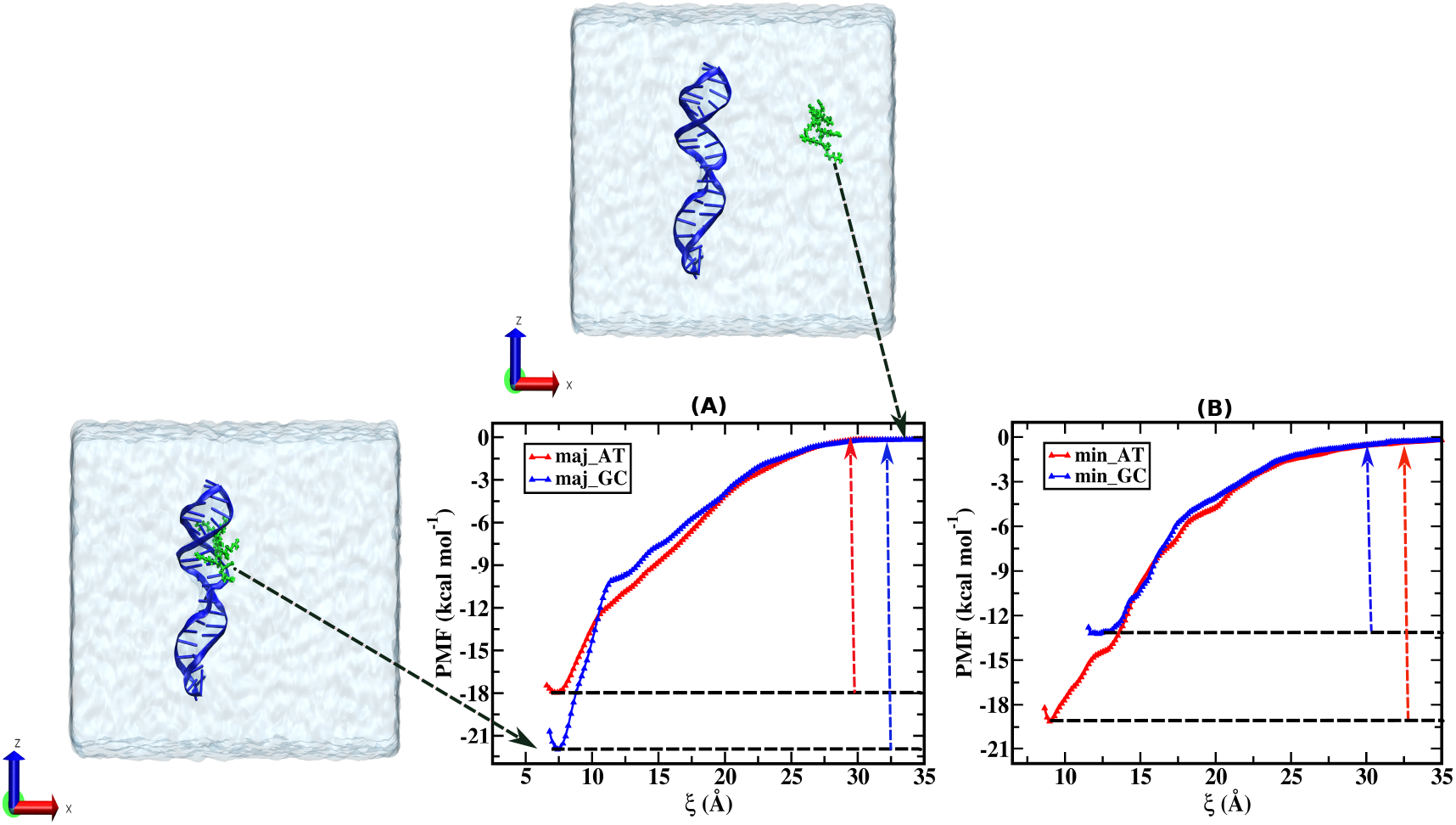
PMF profiles for the Umbrella Sampling of the arginine-rich cationic peptides (protamine-template) along the reaction coordinates from binding regions (range of DNA residues) of major and minor grooves of twenty base pair DNA utilizing the umbrella sampling approach. **[A]** depicts the free energy profiles of peptide dissociation from the major grooves of d(AT) and d(GC) in red and blue color, respectively. **[B]** displays free energy profiles of peptide dissociation from the minor grooves of d(AT) and d(GC) in red and blue color, respectively. The ions are not displayed for clarity in the snapshots of water boxes (in ice-blue color) containing the DNA-peptide complex on the left and top of (A). The conformations of DNA-peptide complexes in the bound and unbound regions are shown in section S.8 in SI.

### E. Explanation of sequence-specific recognition

Sequence-specific recognition during protein binding to the DNA groove is typically accomplished using a combination of direct and indirect readouts. Direct readouts rely on hydrogen bonding and precise stereochemical fitting, whereas indirect readouts depend on DNA’s conformational flexibility^2^. In essence, the protein-DNA interactions and DNA deformation are the two critical mechanisms that play a significant role in the quantitative recognition of a peptide binding site^66^. For polyamine binding to the DNA groove, the experimental Raman-spectra suggests that interaction with negatively charged phosphate groups is strong in minor grooves, and the sequence context of DNA plays a secondary role in recognition^69^. AT-rich regions of DNA have narrower minor grooves and more negative electrostatic potential than GC-rich regions^2^. Using 3dna software^70^, we computed the average minor groove width of the DNA where the peptide is bound. The minor groove width of the d(AT) duplex is 13.38 ± 4.49 Å, while the d(GC) duplex is 10.74 ± 3.71 Å as listed in Table II, due to the stacking interaction of the planar guanidinium side group with the DNA bases at the d(AT) minor groove, causing the structural deformation and minor groove widening.

**TABLE II.** Groove width and bending angles of the DNA duplexes from unbiased simulation.

| Sequence<br>(Minor Groove) | Groove<br>Width<br>(Å) | Bending<br>Angles<br>(°) | Sequence<br>(Major Groove) | Groove<br>Width<br>(Å) | Bending<br>Angle<br>(°) |
| --- | --- | --- | --- | --- | --- |
| **min_AT** | $13.38 \pm 4.49$ | $40.91 \pm 17.52$ | **maj_AT** | $20.37 \pm 7.91$ | $38.17 \pm 13.58$ |
| **min_GC** | $10.74 \pm 3.71$ | $32.04 \pm 10.58$ | **maj_GC** | $19.67 \pm 7.52$ | $73.70 \pm 19.31$ |

For the computation of the groove width and bending angle of the duplexes, we took only the central ten base pairs of the duplexes that include the groove where the peptide is bound.

The 3dna calculates the minor and major groove widths using direct P-P distances; to account for the phosphate groups’ vdW radii and to compare the findings with those obtained using the FreeHelix and Curves software, 5.8 Å must be subtracted from the numbers^71^. Using DNAphi web server^72^, we estimated the average electrostatic potential of the minor groove for the given sequences of DNA at the peptide binding site. We found the electrostatic potential to be-6.79 *kT/e* for the d(AT) duplex and-4.81 *kT/e* for the d(GC) duplex, with *k, T*, and *e* denoting the Boltzmann constant, simulation temperature, electron charge (magnitude), respectively. Peptide binding in the minor groove is characterized by the side chains extending toward the exterior and the peptide backbone toward the groove floor^73^. From these facts, we can say that the narrower minor groove with high negative electrostatic potential provides better stereochemical close-fitting to the peptide with positive guanidium residues of arginine extending towards the negative phosphate residues of the backbone, making the binding of the peptide stronger in the minor groove of d(AT) duplex than in the minor groove of the d(GC) duplex. During DNA-protamine binding, the planar guanidinium side group of arginine (ARG) forms stacking interactions with the DNA bases, causing significant deformation and unwinding of AT-rich DNA. These stacking interactions occur alongside the electrostatic interactions between protamine arginines and DNA phosphates. Therefore, stacking-induced distortion occurs in the minor groove of the AT-rich DNA motif, as evidenced by the highest total RMSD among the four complexes. DNA becomes more flexible by unstacking or unwinding caused by ARGs, and they can significantly reduce intra-DNA(same strand) repulsion between phosphates.

Using 3dna software, we found the bending angle to be 40.91 ± 17.52 ^°^ for d(AT) duplex and 32.04 ± 10.58 ^°^ for d(GC) duplex, for the case of peptide bound to the minor groove as listed in Table II. This explains why the peptide remains closer of the minor groove in the d(AT) duplex, but near the edge of the groove in the d(GC) duplex. Also, the narrow minor groove binding region of d(AT) duplex is more negative than that of d(GC) duplex as shown in Figure 8[B] with three acceptor electron lone pairs(*δ* −), whereas d(GC) has two.^68^. As the backbone of peptide remains more fitted to the minor groove of d(AT) duplex (due to a more straight, extended, and narrow groove) and the positive amide groups of the side chains extend outwards interact more strongly with negative phosphate backbone^74^. This is why the binding of peptides in the minor groove is more favorable in the AT-rich motifs of DNA than in GC-rich motifs. As the d(AT) interacts more closely with the peptide than d(GC) through intercalation in the minor groove, the stereochemical fitting of the peptide prefers d(AT) *>* d(GC) for the sequences taken in this work. Hence, the preference of binding in the minor groove by the peptide follows d(AT) *>* d(GC). That means AT-rich motifs are highly preferred, and GC-rich motifs are least preferred by the cationic peptide for binding in the minor groove.

**FIG. 8.**
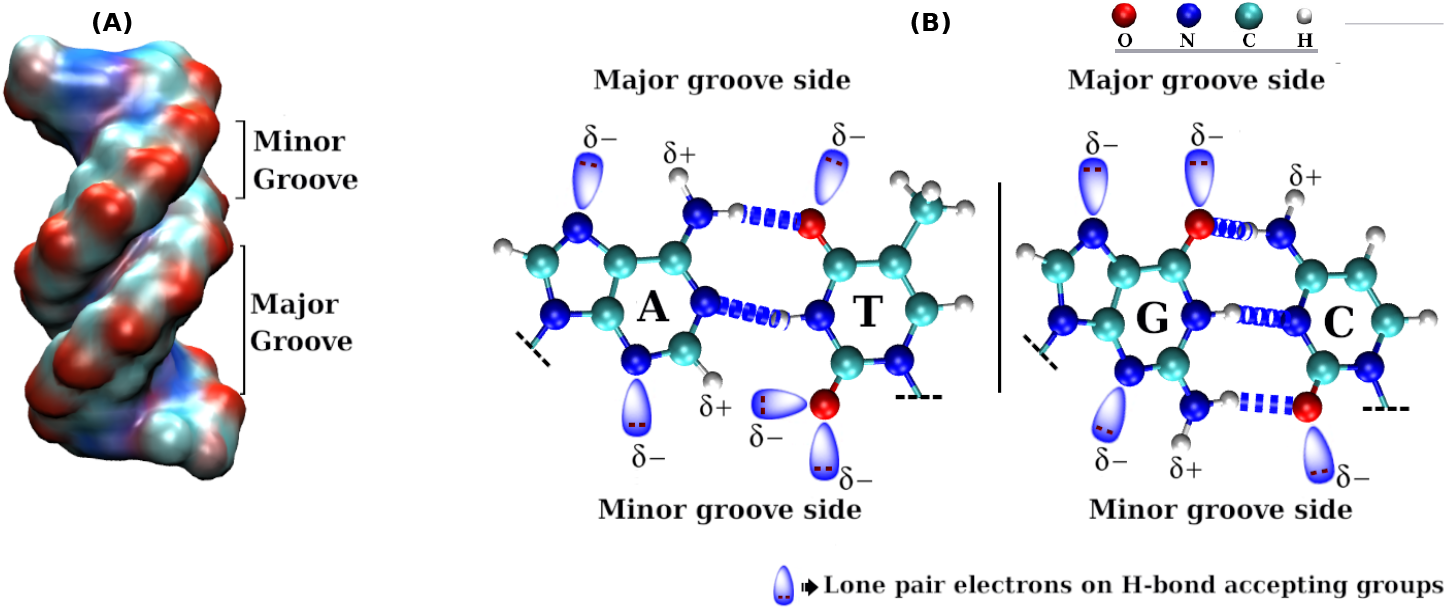
Major and minor grooves of the DNA duplex and the donors and acceptors for H-bond formation: **[A]** schematic of the wider major groove as well as narrower minor grooves of the DNA and **[B]** donors (*δ* +) and acceptors (*δ*−) presented by the A-T and G-C base pairs towards the Major(top) and Minor(bottom) grooves of the DNA. The presence of methyl group (CH_3_) of thymine limits the amount of H-bond acceptor atoms for the formation of H-bonds towards the major groove side of the A-T base pair^67^. Also, the minor groove floor of the A-T base pair is much more negative with three acceptor electron lone pairs(*δ* −) than that of the G-C base pair with two acceptor lone pairs (*δ* −)^68^.

For major groove binding, the direct contacts (H-bonds and other contacts) and the DNA shape modulation play a role in the binding affinity^75^. From earlier sections, we found that the H-bonds, the number of contacts, and different interaction energies between the peptide and the DNA major groove showed the preference order d(GC) *>* d(AT). During major groove binding, the DNA bends over the peptide to enhance the interaction between the peptide and the DNA, encasing the peptide in a deepened, constricted, and moderately electronegative major groove^74^. The bending of the DNA towards the major groove makes it slightly constricted and deeper, enhancing its electrostatic potential. Using 3dna software, we found the bending angle to be 38.17 ± 13.58^°^ for d(AT) duplex and 73.70 ± 19.31^°^ for d(GC) duplex, for the case of peptide bound to the major groove. It shows that shape modulation is more significant on d(GC) duplexes than d(AT) duplexes. DNA shape modification through helix bending or other deformation facilitates the emergence of a series of hydrogen bonds or non-polar interactions between the peptide and DNA, which would be considerably less common in the absence of the deformation^75^. The stiffer nature of d(AT) duplex should therefore be one of the prime cause to have least binding affinity of the peptide in the major groove of the d(AT) duplex. The reason for the high preference for d(GC) rather than d(AT) for major groove binding remains in the fact that arginine prefers guanine(G) bases most prominently for binding^76^. The computation of interaction strength between serine and various nucleobases, as well as the related binding energies, revealed an energetic ordering of guanine *>* cytosine *>* thymine *>* adenine *>* uracil, based on the first-principles analysis^77^. Also, the peptides are sterically blocked by the methyl groups of thymine(T) bases^67^ for AT-rich motifs. That means the clustering of methyl groups in AT-rich DNA sequences repels the peptide from the major groove and hinders its effective binding to the groove^67,78^. Therefore, the lower preference of the peptide to be attached in the major grooves of d(AT) duplexes compared to the d(GC) duplex can be explained by the existence of the methyl group of thymine, which limits the number of H-bond acceptor atoms allowing the formation of H-bonds between peptide and DNA bases.

These results substantiate our anticipation that the highly extended geometry of the peptide permits more area for effective interaction with the DNA minor groove by aligning its backbone in the groove, extending the side chains outward, and thereby increasing the number of contacts with the DNA. The results of this work are consistent with those of an X-ray diffraction analysis^21^, which shows the number of contacts made by the peptide with DNA when adsorbed in the minor groove is more than the expected one. The higher number of contacts found in minor groove binding should be associated with the groove’s intercalative interaction (see section S.10 in SI for the qualitative estimation of stacking/intercalation for the poly(AT) minor groove binding) with guanidinium residues of arginine. Because the DNA minor groove is generally too small to allow peptide structural components, therefore we have found some structural distortions on the backbones of DNA duplexes and their Watson-Crick base pairs during the binding of the peptide in the minor grooves of d(AT)^30^.

## IV. CONCLUSION

Protamines are basic proteins that cause sperm cells’ DNA to be securely coiled and compressed to protect the genetic material. Therefore, a thorough understanding of the preferred DNA sequence and grooves for the protamine binding is extremely important to uncover. We have explored the sequence-specific (AT vs. GC) binding affinity of an arginine-rich short-cationic peptide in the major and minor grooves of DNA through atomistic MD simulation. We found that the peptide exhibits a smaller RMSD when attached to the major and minor grooves of d(GC) compared to d(AT). The peptide is found to be adhered to the backbone of DNA rather than being fitted inside the groove in case of minor groove binding of d(GC). The COM distance between the peptide and the DNA binding site is lower for a peptide that fits properly in the grooves of the DNA. We found that when the peptide is in the major groove of the d(AT) duplex, the COM distance is larger compared to that bound at the d(GC) duplex. For minor groove binding, the peptide is located more profoundly in the minor groove of the d(AT) duplex than in the d(GC) duplex. The structural analysis shows that, when it comes to minor groove binding, the peptide fits better in the minor groove of d(AT) than d(GC), whereas it binds strongly to the major groove of d(GC) than to d(AT).

H-bonds play a major role in DNA-protein interactions, allowing the protein to directly read the DNA. The peptide forms more H-bonds with the major groove of d(GC) than it does with d(AT). Conversely, d(AT) exhibits more H-bonds than d(GC) for minor groove binding. The total contact numbers between the peptide and DNA grooves show a similar tendency. The NB interactions (both vdW and electrostatic) are critical for peptide-DNA complex stability. For major groove binding, d(GC) has a higher total NB energy than d(AT), but for minor groove binding, d(AT) outperforms d(GC) due to stacking interactions.

The MMGBSA binding free energy calculations confirm that the peptide prefers d(GC) in the major groove, while in the minor groove, it prefers d(AT), and the interactions are enthalpy-driven. The PMF profiles using Jarzynski’s Equality also support the binding preferences of the peptide in major and minor grooves of d(AT) and d(GC), the same as MMG-BSA calculation. Our advanced sampling techniques, such as umbrella sampling, also reveal protamine’s preference for d(GC) in the major groove and d(AT) in the minor groove, which is consistent with all previous structural and thermodynamic insights.

The bending angle measurement revealed that d(GC) bends more than d(AT) towards the major groove (Table II). As a result, the stiffer nature of the d(AT) duplex should be one of the primary reasons for the peptide’s low binding affinity in the major groove of the d(AT). In addition, AT-rich sequences have a structurally narrower, electrostatically more negative minor groove, which provides stereochemical close-fitting to the peptide for binding in the minor groove of d(AT). So, the better the stereochemical close-fitting of the peptide and stacking interaction with DNA bases, the stronger the peptide binding in the minor groove of the d(AT) duplex, resulting in some deformation of the DNA helix. These findings suggest that the arginine-rich cationic peptide prefers AT-rich motifs for the minor groove and GC-rich motifs for the major groove bindings.

Therefore, through the microscopic analysis of the simulation trajectories, we found that the arginine-rich cationic protamine-template shows a sequence-specific preference for binding in the major and minor grooves of DNA. Furthermore, the entire behavior of the actual long protamine may not be reflected in the peptide sequence used as a template. Further studies containing the complete protamine structures are needed.

## V. CONFLICTS OF INTEREST

There are no conflicts to declare.

## ACKNOWLEDGMENTS

S.M acknowledges the SRF fellowship from CSIR, India. Y.H.J. and Y.L. thank the National Research Foundation of Korea (2019R1A2C2003118 and 2021H1D3A2A01099453) for financial support. We thank SERB, India for financial support through CRG/2021/003659

## DATA AVAILABILITY STATEMENT

The data validating the findings of our simulation study are available upon request to the corresponding author.

## Notes

### Competing Interest Statement

The authors have declared no competing interest.

